# Expression Profile of Primary Microcephaly Genes *Cdk5rap2*, *Cep63* and Their Paralogues *Pde4dip*, *Ccdc67* in Mouse Embryonic and Adult Tissues

**DOI:** 10.1101/2023.12.16.571987

**Authors:** M. Khurshid, K. Fatima, T. Azam

## Abstract

Autosomal recessive primary microcephaly (MCPH) is characterized by the prenatal reduction in the human brain growth, without any change in the cerebral architecture. MCPH can result from the bi-allelic mutations in at least thirty genes and still counting on. MCPH genes majorly code for a centrosomal protein subset, cell cycle proteins, nuclear proteins and cytoskeletal proteins. Centrosomal subset is the most highly characterized subset of MCPH proteins. *Cep63* and *Cdk5rap2* with their paralogues *Ccdc67* and *Pde4dip* play important role for the normal assembly of spindles and in DNA damage response. They regulate neuroepithelial cell division in the developing brain, so decrease in centrosome number results in genomic instability. Study presented in this manuscript was aimed to identify the tissue specific expression of genes with their paralogues (*Cep63* and *Ccdc67*) and (*Cdk5rap2* and *Pde4dip*) in developing and adult normal mice tissue cells. For this purpose we successfully generated timed pregnant mice at the peak of neurogenesis and 36 samples were screened for the expression of these genes and their paralogues. RNA was extracted from the mice tissues using trizol reagent. Extracted RNA was reverse transcribed and cDNA was subjected to PCR amplification using gene specific primers, generated by using primer3 at the intron-exon junction to avoid the genomic interference. One house keeping gene β-actin was used as a positive control. This study reveals that *Cep63 and Ccdc67* paralogues are playing a suggestive collective role in the mice development as shown by multiple length bands. *Cdk5rap2* an MCPH gene and its paralogue *Pde4dip* band suggested that maybe they might have redundant roles in body organs apart from the brain. These proteins have tissue-specific expression in all studied tissues from the developing and adult mice. We identified multiple bands in the different samples of all four genes which might reflect towards multiple transcript variants.

## Introduction

Autosomal recessive primary microcephaly (MCPH) is infrequent, hereditary and neurogenic Mendelian disorder. In this disorder the size of cerebral cortex is reduced, and as a result, the head circumference is significantly decreased [3-5]. When the circumference of head is >3 SD [6] below the mean for the gender and age matched controls termed as microcephaly [7]. Microcephaly is strongly associated with mental illness [8]. There is no well-defined classification of microcephaly but generally it has two types. When the brain fails to achieve the normal brain size during pregnancy it is called Prenatal/Congenital or Primary Microcephaly, but when the brain size is normal during childbirth but does not grow properly after that it is called Postnatal/Acquired or Secondary Microcephaly [4, 9-11]. Range for the occurrence of microcephaly is from 1.3 to 150 per 100,000 depending upon the range of SD and type of population. MCPH is more common in Asians and Arabs (consanguineous) with incidence of 1/10,000 which is more than in White (non-consanguineous) [7].

Genetic and non-genetic conditions that may underlie microcephaly are genetic syndromes, metabolic conditions (e.g., maternal phenylketonuria), exposure to radiations and teratogenic agents, deprived fetal care, in-utero infections, cerebral anoxia, severe mal nourishment, alcohol consumption by mother during pregnancy and non-accidental head injury [9, 12].

Affected individuals with MCPH have lessened cerebral cortical volume with simplified gyral pattern. Patients of MCPH can have mental retardation ranging from mild to moderate with non-progressive intellectual impairment, defective cognitive abilities, and facial distortions including sloping forehead which is not always present, short stature, hyperactivity, stroke/convulsions/sudden illness, complications in coordination, maintaining balance and other neurological deformities/defects of brain. Majority of MCPH patients can be easily handled and can acquire self-help expertise [7, 8, 13].

Neuronal migration and neural apoptosis are the major contributors of MCPH disorder. MCPH genes have higher expression in the neuroepithelium. In neuroepithelium, Neuronal progenitors (neuroepithelial cells NEP and radial glial cells RG) are the chief cells and majority of neurons originate from them. In the developing neuroepithelium nerve cells undergo specialized form of mitosis. In **symmetric proliferative mitosis** pattern of mitosis, two neural progenitor cells are formed from the progenitor and this is due to mitotic spindle being perpendicular to the neuroepithelium, while in **asymmetric** divisions, one progenitor and one post mitotic neuron is formed that is in the plane of neuroepithelium **[3, 14, 15]** (Figure 1). During development, division type and timings decides the final number of neurons. Cortical disorders may results because of abnormal regulation of these divisions like microcephaly and microlissencephaly [16]. Cerebral cortex controls the processes of communication, creativity and rapid decisions etc. However, poor growth of brain in microcephaly is the results from abnormality in this pathway by any of the involved gene [17].

The inherited pattern of MCPH is autosomal recessive, which means that both copies of the gene in each cell carry mutations. People with MCPH receive one mutated copy of the gene from each parent, but the parents commonly don’t exhibit any symptoms of this condition [18]. During cortical neurogenesis mostly all these genes are expressed in the essential germinal zone in the cerebral cortex, called the ventricular zone (VZ), which plays a part in multiplication (proliferation) of neural progenitor cells (NPCs). Intriguingly, the centrosome has been involved in the pathogenesis of several MCPH disorders [12]. Thirty causative loci (MCPH1-MCPH30) have been identified, responsible for causing microcephaly and still progressing **[12, 19]**.

### Cdk5rap2

The basis of MCPH3 is homozygous mutations of cyclin-dependent kinase 5 regulatory Subunit-associated protein-2 *(Cdk5rap2)* [20]. There are 38 exons in the gene of 191290bps length with 5862 ORF encoding 1893 amino acids in this protein. *Cdk5rap2*, present on chromosome 9q33.2, is a centrosome-based protein, and mRNA for this protein is broadly expressed in embryonic tissues of mouse and human. Cohesion and condensation processes of chromosomes are carried out by two ATPase dependent domains of *Cdk5rap2* protein [11, 20]. Spindle checkpoint regulation is controlled by *Cdk5rap2*. Any loss of *Cdk5rap2* results in chromosomal mis-segregation and decreased protein expression of spindle check point. During neuronal progenitor cell division, premature transition of cells from symmetric to asymmetric is a result of Cdk5rap2 mutation. This mutation results in dysfunctioning of centrosome, reduced rate of cell survival and ultimately reduction in progenitor pool [18]. *Cdk5rap2*mutations disturb the assembly of mitotic spindles, thus this affects the neurogenic mitosis [7]. During early neurogenesis, *Cdk5rap2* has higher expression in the spinal cord and neuroepithelium of brain. In mouse *Cdk5rap2*protein is localized in pericentriolar material all through the stages of cell cycle [7].

### Pde4dip

*Pde4dip* (Myomegalin) is a homologous paralogue of *Cdk5rap2* in mammals. *Pde4dip*, a centrosomal protein, has a predominantly complementary expression pattern to *Cdk5rap2*. During neurogenesis both these proteins have functional homology in their roles. So this increases the microtubules production in centrosome and apply a negative control over their inhibitors [21].

### Cep63

It is centrosomal protein 63 and its gene is present on chromosome no. 3q22.2. *Cep63* was first identified by using mass spectrometry, as it is an active component of purified centrosomes and has two domains. In human cells and in mouse cells *Cep63* forms interaction with *Cep152* and plays a key role in centriole duplication. So reduction in the level of *Cep63* and *Cep152* at centrioles results in the reduction in pericentriolar material (PCM) size, and result in inefficient duplication of centrioles [22]. Any mutation in *CEP63* results in Seckel syndrome in humans. It is characterized by the dwarfism and microcephaly, a human disease [23]. Mutation in *Cep63* is homozygous in nature and causes MCPH. *Cep63* forms a complex with *Cep152*, which act as a conserved factor for centrosome duplication. Both of these proteins are important for the maintenance of centrosomal number at a normal level. Impairment caused by this mutation affects the growth of cerebral cortex [24]. Mutation in *Cep63* is homozygous in nature and causes MCPH. *Cep63* forms a complex with *Cep152*, which act as a conserved factor for centrosome duplication. Both of these proteins are important for the maintenance of centrosomal number at a normal level. Impairment caused by this mutation affects the growth of cerebral cortex [24].

### Ccdc67

In invertebrates *Cep63* has a paralogue called as *Deup 1* (deuterosome protein 1, also known as *Ccdc67*). Both of these proteins have 37% identity in mouse. Epithelial cells of murine trachea and multi-cilia-abundant tissues are highly enriched in messenger RNA of *Ccdc67*.

The brain in humans gives increased behavioral and rational skills leading to human inventiveness. The human brain is behind our cognitive capacities which leads to storage of the memory, our attention gestures, language deriving, learning skills and consciousness[25].

Mice are used as a model to study human diseases owing to their genetic manipulation ability. Among animal models, mice are the most suitable and handy ones with complete genome sequences. The evolutionary divergence of 19 million years is present between the mice and human with comparable brain organization [26, 27].

## Material and Methods

**Figure 1:**
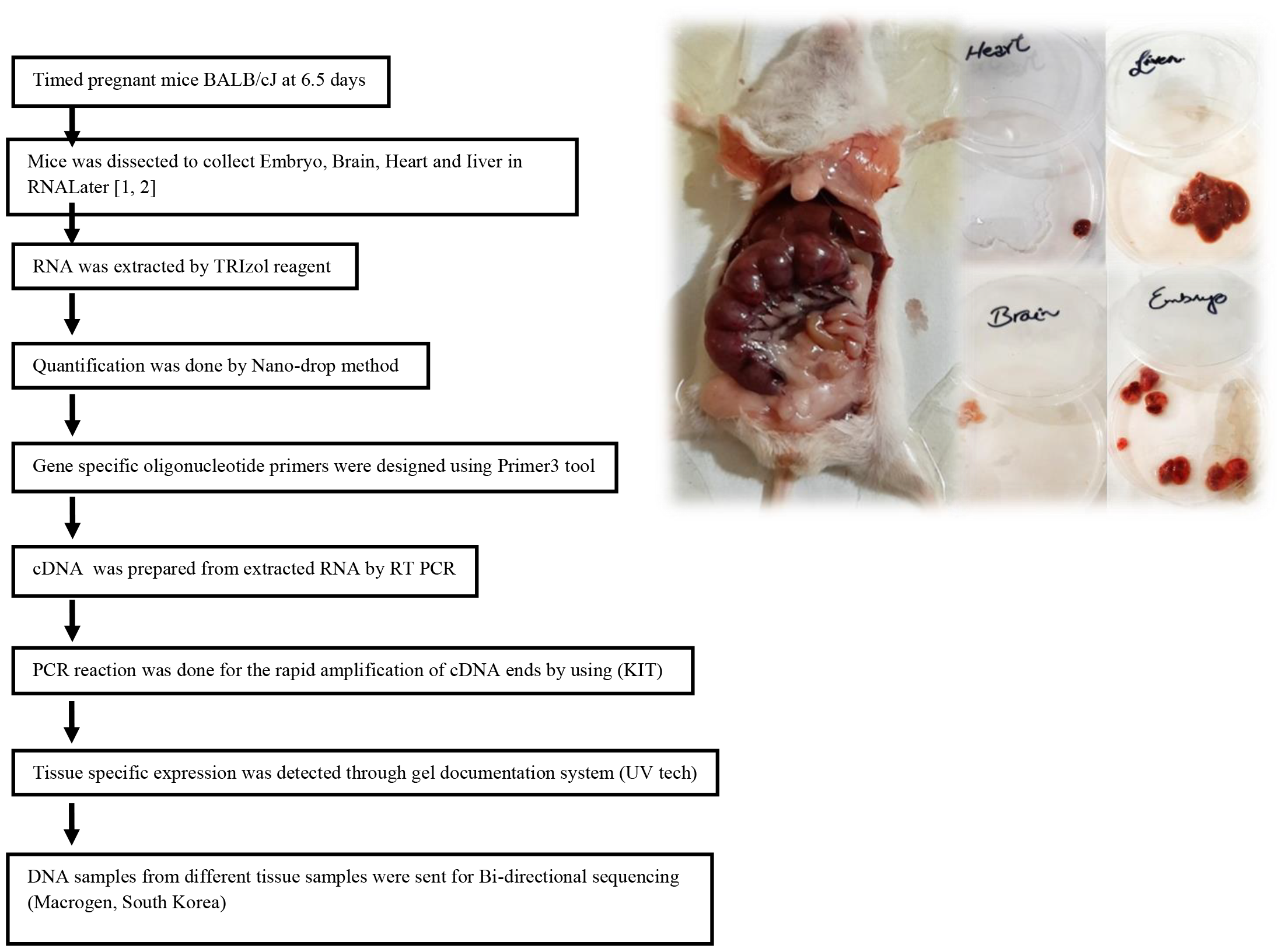
Full view of mice dissection and separated organs.

## Primer Designing

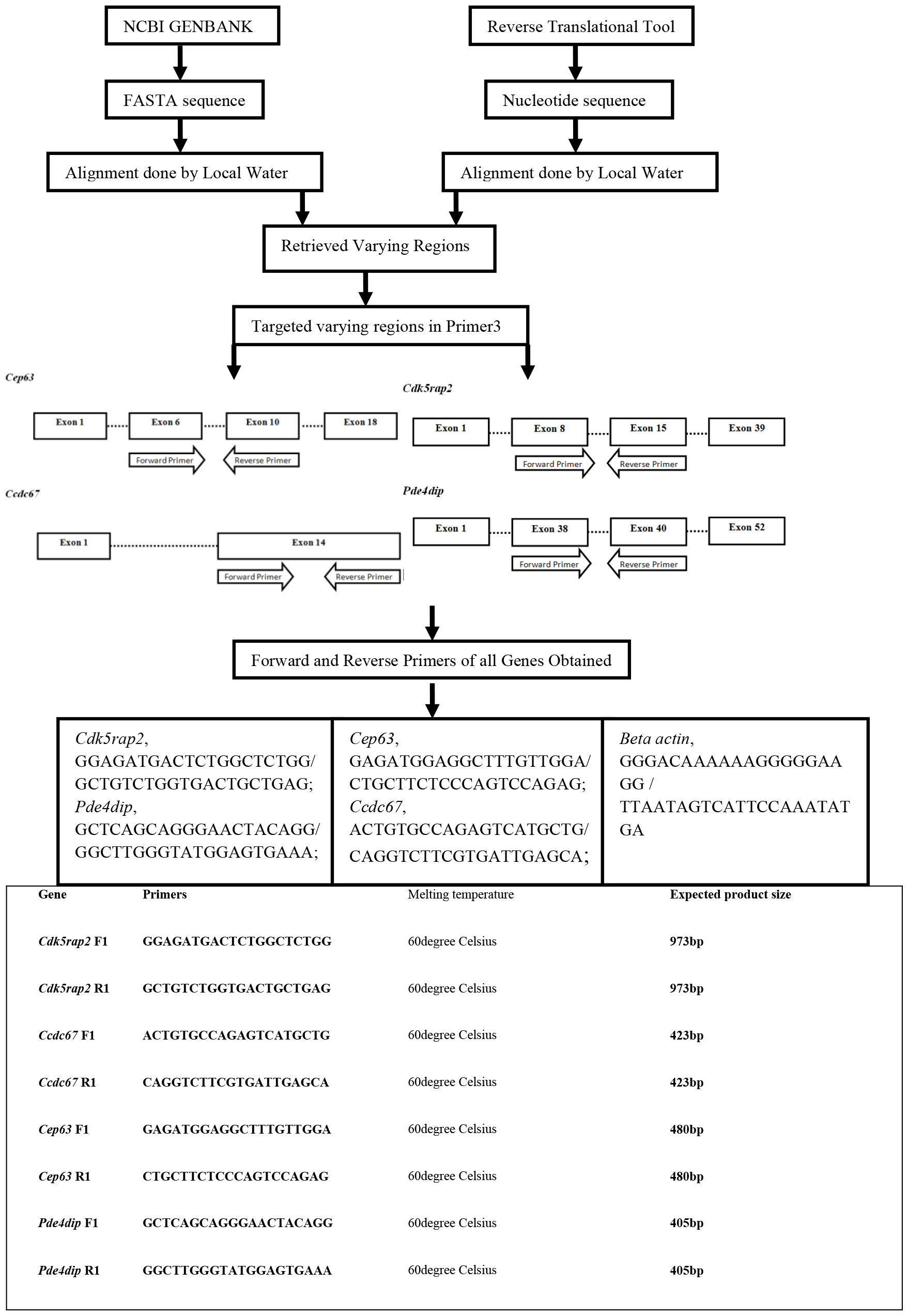

## Results

RNA was extracted from the tissue samples of pregnant mice. Total 4 samples including embryo, adult brain, adult heart, and adult liver, were analyzed for the expression of the selected MCPH genes and their paralogues. RNA was extracted using TRIzol reagent (Thermofisher scientific). RNA was quantified by a Nano-drop method. The quality of extracted RNA was determined by absorbance of RNA solution that was measured at 260 nm as well as at 280 nm. RNA quantification data of all samples has been summarized in the following table:

The first strand cDNA was developed using Poly-dT primer and gene specific primers were used to make double stranded DNA. Expression of the selected MCPH genes and their respective paralogue was documented on the gel shown in Table: 2 and Table: 3.

**Table 1:** RNA Quantification.

| Sample ID |  | A260 | A280 | 260/280 | Concentration ng/μl |
| --- | --- | --- | --- | --- | --- |
| Brain | B1 | 0.594 | 0.318 | 1.867 | 475.32 |
|  | B2 | 0.806 | 0.635 | 1.269 | 644.853 |
| Embryo | E1 | 2.406 | 1.224 | 1.966 | 1924.52 |
|  | E2 | 1.067 | 0.885 | 1.205 | 853.493 |
| Heart | H1 | 0.69 | 0.393 | 1.754 | 552.12 |
|  | H2 | 2.214 | 1.585 | 1.397 | 1771.493 |
| Liver | L1 | 3.362 | 2.284 | 1.472 | 2689.96 |
|  | L2 | 3.19 | 2.847 | 1.121 | 2552.373 |

**Table 2:** Expression analysis of *Cep63* and its paralogue *Ccdc67* in mice tissue samples by using primer on the Gel.

| Sample name | Marker | Brain 1 | Brain 2 | Embryo 1 | Embryo 2 | Heart 1 | Heart 2 | Liver 1 | Liver 2 |
| --- | --- | --- | --- | --- | --- | --- | --- | --- | --- |
| bp |  |  |  |  |  |  |  |  |  |
| Sample name | Marker | Brain 1 | Brain 2 | Embryo 1 | Embryo 2 | Heart | Liver |  |  |
| bp |  |  |  |  |  |  |  |  |  |

**Table 3:**
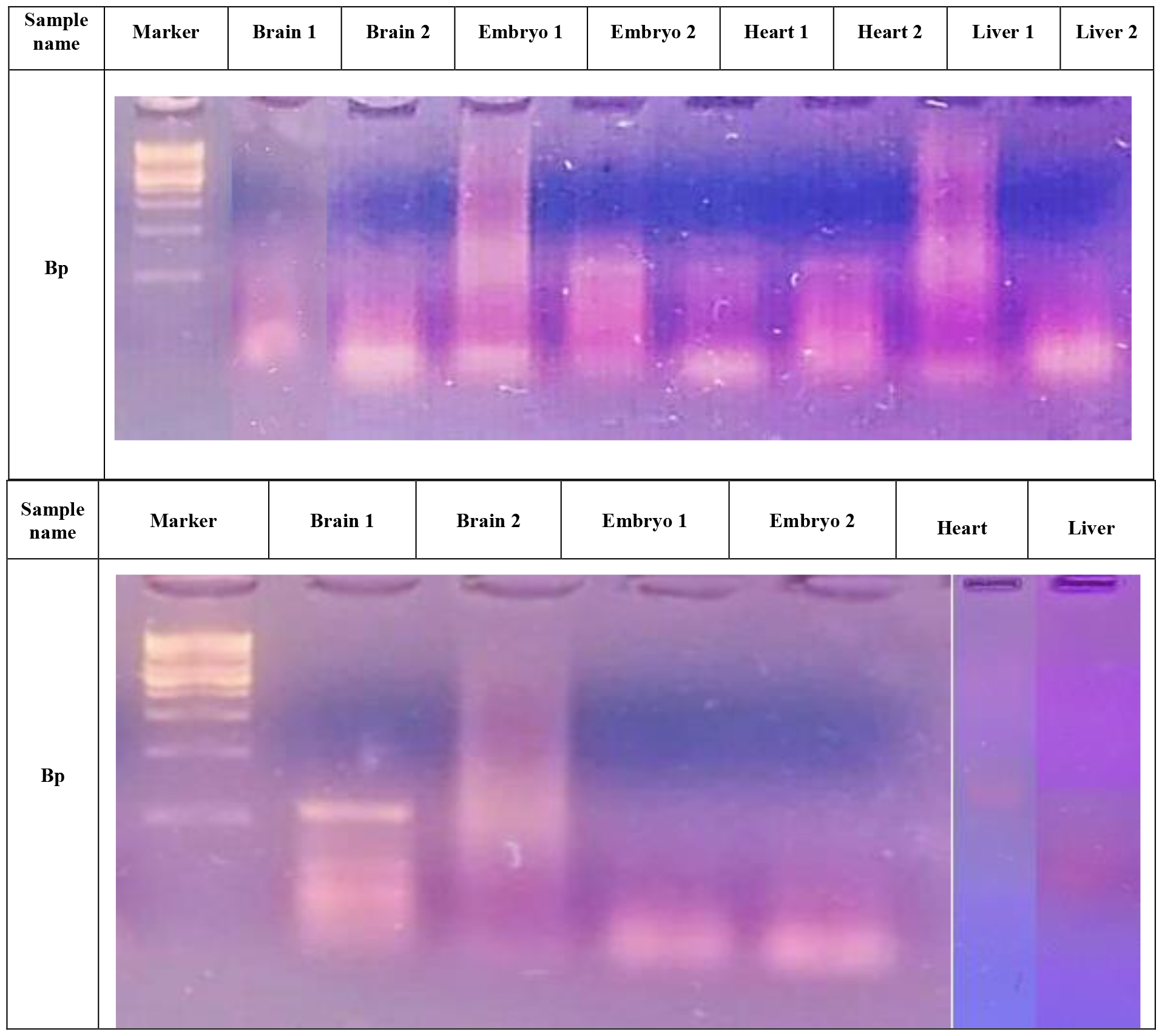
Expression analysis of *Cdk5rap2* and its paralogue *Pde4dip* in mice tissue sample by using primes on Gel.

The control used was for β-actin which is a housekeeping gene shown in Table: 4.

**Table 4:**
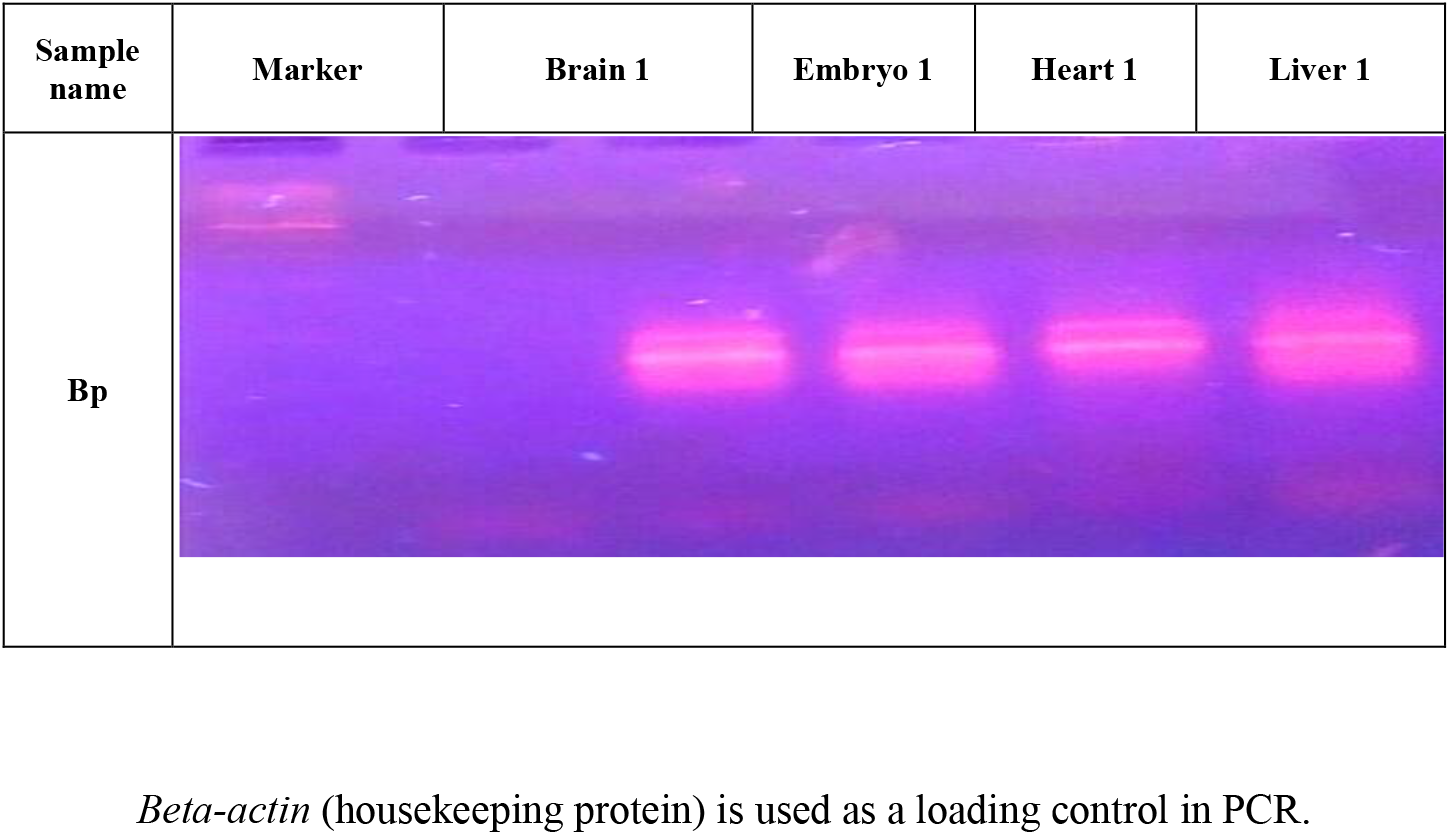
Expression analysis of *β-actin* in mice tissue samples by using primer on the Gel.

| Sample name | Marker | Brain 1 | Embryo 1 | Heart 1 | Liver 1 |
| --- | --- | --- | --- | --- | --- |
| Bp |  |  |  |  |  |

*Beta-actin* (housekeeping protein) is used as a loading control in PCR.

In *Cep63*, heart brain and embryo has two bands of 650 bp and 480bp while in liver a single band of 480 bp appears. In heart and brain band of 480 bp is of higher intensity than of 650 bp while in embryo the band of 650 bp is more intense than 480 bp. In *Ccdc67*, heart and liver shows a single band of 423 bp while brain and embryo have two bands of 970 bp and 423 bp. The band of 970 bp in embryo and brain is of higher intensity than 423 bp.

In Cdk5rap2, heart, liver and embryo have two bands of 973 bp and 408 bp. The band of 408bp of heart is of higher intensity than of 973 bp while in liver and embryo both bands are of same intensity while brain has only one band of 408 bp. In Pde4dip, brain is showing 3 bands of 1050 bp, 550 bp and 405 bp having different intensity suggesting the presence of multiple isoforms in cerebral cortex while heart, liver and embryo have a single band of 405 bp.

## Discussion

It was an arduous task to generate timed pregnant mice at the peak of neurogenesis. The estrous cycle was tracked by examining the vaginal opening to make an initial observation and further verified by vaginal cytology. At the Estrous stage two female mice were housed overnight with one male per cage as ovulation occurs at this stage.

The very next early morning, the presence of plug in vagina indicates that copulation has occurred which may lead to pregnancy. After the considerable trouble we attained the aim of our study and got mice pregnant embryos at E6.5 and E14.5 (The results in this paper refer to both embryo stages). The subject female mice was dissected and adult organs of the heart, brain, liver and embryos were collected in falcon tubes containing RNA stabilizing reagent to protect and freeze the samples for later processing from pregnant dam.

RNA was extracted from collected mice tissues followed by the quantification of RNA by Nano-drop method. Quantity of RNA in each sample was given by Gen5 software while the quality of extracted RNA was measured by its absorbance at 260 nm in addition to 280 nm. As the ratio of A260/A280 was between 1.0-2.0 so these samples were further processed for cDNA preparation.

Primers were generated by using primer3 at the intron-exon junction to avoid the genomic interference. PCR, an invitro amplification process, was performed at specific conditions for amplification of genes and their paralogues in order to get the PCR products. The resulted products of PCR amplification along with a ladder and negative control were analyzed at 2% agarose gel. The gel was run at 70 Volts for approximate time and documented through a gel documentation system (UV tech). Gel Images were taken and saved for future use.

This study reveals that *Cep63/ Cdk5rap2* and their paralogues may have a complex expression in tissues with more than one isoforms/splice variants. They paralogues might be playing a redundant role to the MCPH genes in the body of the affected individual which is all functional apart from small cerebral cortex. One or all of the isoforms/splice variants maybe specific for functional embryonic brain (cerebral cortex). Although the MCPH genes are showing more than one isoform which makes things more complex than suggesting that certain paralogue isoform may compensate for its expression in developing cerebral cortex. Here in our results *Ccdc67* is showing two bands in the adult mouse cerebral cortex and two bands in the embryo while *Pde4dip* is showing three bands in adult mouse cerebral cortex but only single isoform in the embryo. Hence to further prove the differential expression of the genes cloning must be done of all the multiple isoforms and translational work in cells and mice must be done. The knockout of different isoforms of both MCPH genes and their paralogues must be done to get functional insight. Cep63 and Cdk5rap2 proteins have tissue-specific expression in all tissues we studied from the developing and adult mice. In the cerebral cortex and embryo brain, there are multiple isoforms these might reflect brain-specific expression. These proteins are highly expressed in mice brain especially developing the cerebral cortex and are highly conserved in evolution so they might play a role in evolution as well.

Moreover, both genes were studied for the isoforms using bioinformatics tools as EMBL-EBI, Ensemble and the resultant proteins of these isoforms were studied for the 3D structures by using PHYRE2 and EXPASY tools.

Different variants were identified for Cdk5rap2 and *Pde4dip* in the mouse tissues which might be behind normal organs of MCPH subjects.. There are two isoforms of *Cep63* in the embryonic brain and adult brain which might hint towards a neurogenesis specific role in the development of some isoform. A heart-specific isoform for *Ccdc67* was identified which at the time was novel and may play a complementary role in development and their combined role might be behind Seckel syndrome. These proteins are highly expressed in mice brain especially developing cerebral cortex and are highly conserved in evolution so they might play a role in evolution as well.

Future work will make clear the genetic players behind mysterious brain disorders such as primary microcephaly.

## Acknowledgements

I want to acknowledge my team for their work. I further extended this work in the form of research project and I thank HEC, Pakistan for providing funds for the project. I want to especially thank my PhD supervisor, Professor C. Geoffrey Woods for the initial work done on the microcephaly genes during my work in PhD.

## Supplemental Data

### Sequencing data for *CCDC67*

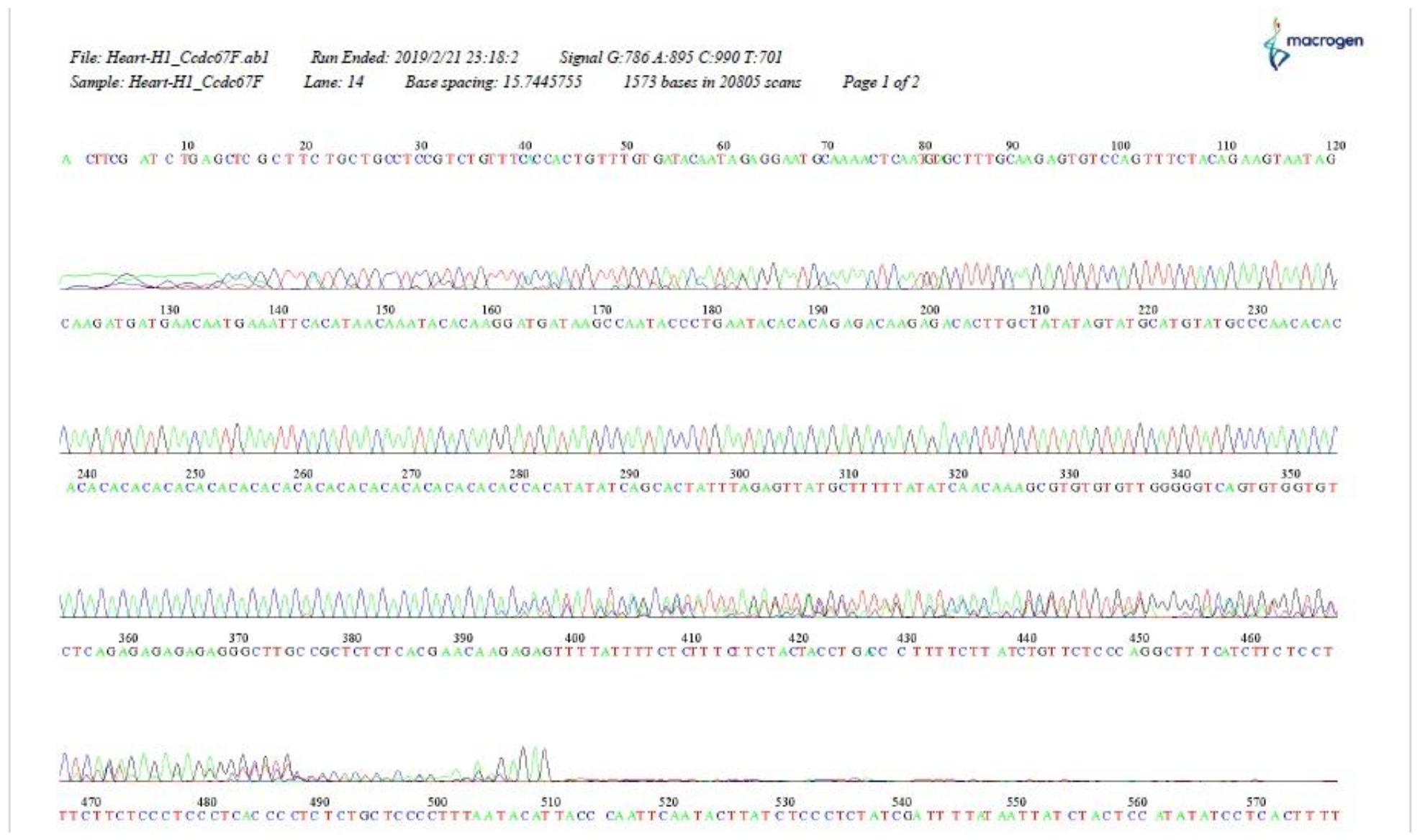

**>190221-R02_C03_Heart-H1_Ccdc67F.ab1 1573**

**ACTTCGATCTGAGCTCGCTTCTGCTGCCTCCGTCTGTTTCACCACTGTTTGTGATACAATAGAGGAATGCAAAACTCAATGTAGCTTTGCAAGAGTGTCCAGTTTCTACAGAAGTAATAGCAAGATGATGAACAATGAAATTCACATAACAAATACACAAGGATGATAAGCCAATACCCTGAATACACACAGAGACAAGAGACACTTGCTATATAGTATGCATGTATGCCCAACACACACACACACACACACACACACA CACACACACACACACACACACCACATATATCAGCACTATTTAGAGTTATGCTTTTTATATCAACAAAGCGTGTGTGTTGGGGGTCAGTGTGGTGTCTCAGAGAGAGAGAGGGCTTGCCGCTCTCTCACG AACAAGAGAGTTTTATTTTCTCTTTCTTCTACTACCTGACCCTTTTCTTATCTGTTCTCCCAGGCTTTCATCTTCTCCTTTCTTCTCCCTCCCTCACCCCTCTCTGCTCCCCTTTAATACATTACCCAATTCAATACTTATCTCCCTCTATCGATTTTATAATTATCTACTCCATATATCCTCACTTTTACTCACTTCTAACCATCCTCAACGACTATACCTTCACCATTTCACCTCCTATCATTTAAGATTACCCGATGAACTTAAACATTTTCATACCCCAGAGAAAAGATTTGTCTTTTCACCTCCTTCTCTATTCGTTCTTTTCCTCTCTCCCCAAAATAGACATATAAACATTTTTTTGCGCGTCTTTCTAGTTCCCATCGACTAGTCCTTTATTATTTTTTTTTTCAAGTTCCTATTTCCGATATGTTAATTTAGTGAATTGGTTATCAGAGTGTTCTATCTGATTTTCCAGGATTAACGTCCTCTGCCACTCTATATTACTTCCTCGTTTATTAGAAGAGGGGCTGTTAGGGGCGCCCCCTTGTTTTATTATATCTTCTCCCCGAACCCATTACGTTAATTTGAGGGTTTTTTTTTAATGTCAATCCTATTGTTTTTATTATTTTACTATCCGGGCTAGTGTCGGGAACAATATTAGCTCTGAATAGTACCTATTCCCTCCTAGCTACACCGGTTTTTTTGCGAAATTTTTTTATAGAGTACTTCTCCCGTACATATTAATTAGTATTCCGATACTTTTTTTATTCCTCACCGAGCACCCGCTACGCGACTTTTTTTTTCTTGCGCTTCTTAAATCTTAATTTTGTGTCGGATATTTATTACCCCCCTCGCCGCCCCCGCATGGCACTATTATATACCCAAATTTTGTTTTTGTGTAGTAATTTTTACTGTTATTTTTTTTACTCCGCACACCAACATTGTTACTTTTTTTGTGCCACATCGTGTACGTACTTTATTATTATTTTTACTCGACGCCCTGGCACCGAATTACAATCGCCCCTTGCTCTGTATATTATCTGCATGAGACTTTTGTAAATACTATTACAGTAATATTATTTTGTGTGAGGGTAGCTCCGACGACAGCGCATGAATAATATAATGTTTCAGACAATTGAGGCAACCCATTGTGCACATGGTTGAATTCTAGGGTCCTACCGC**

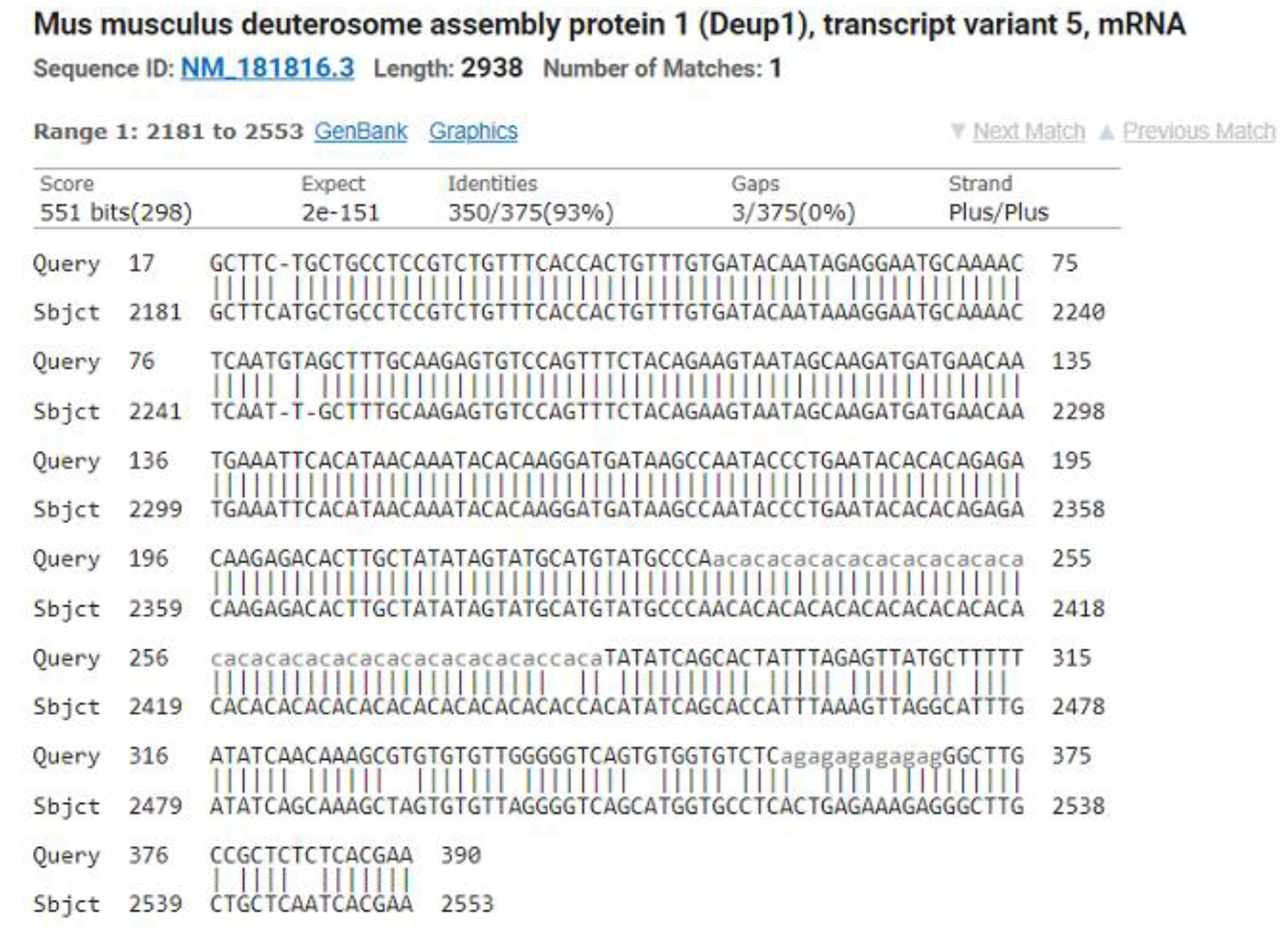

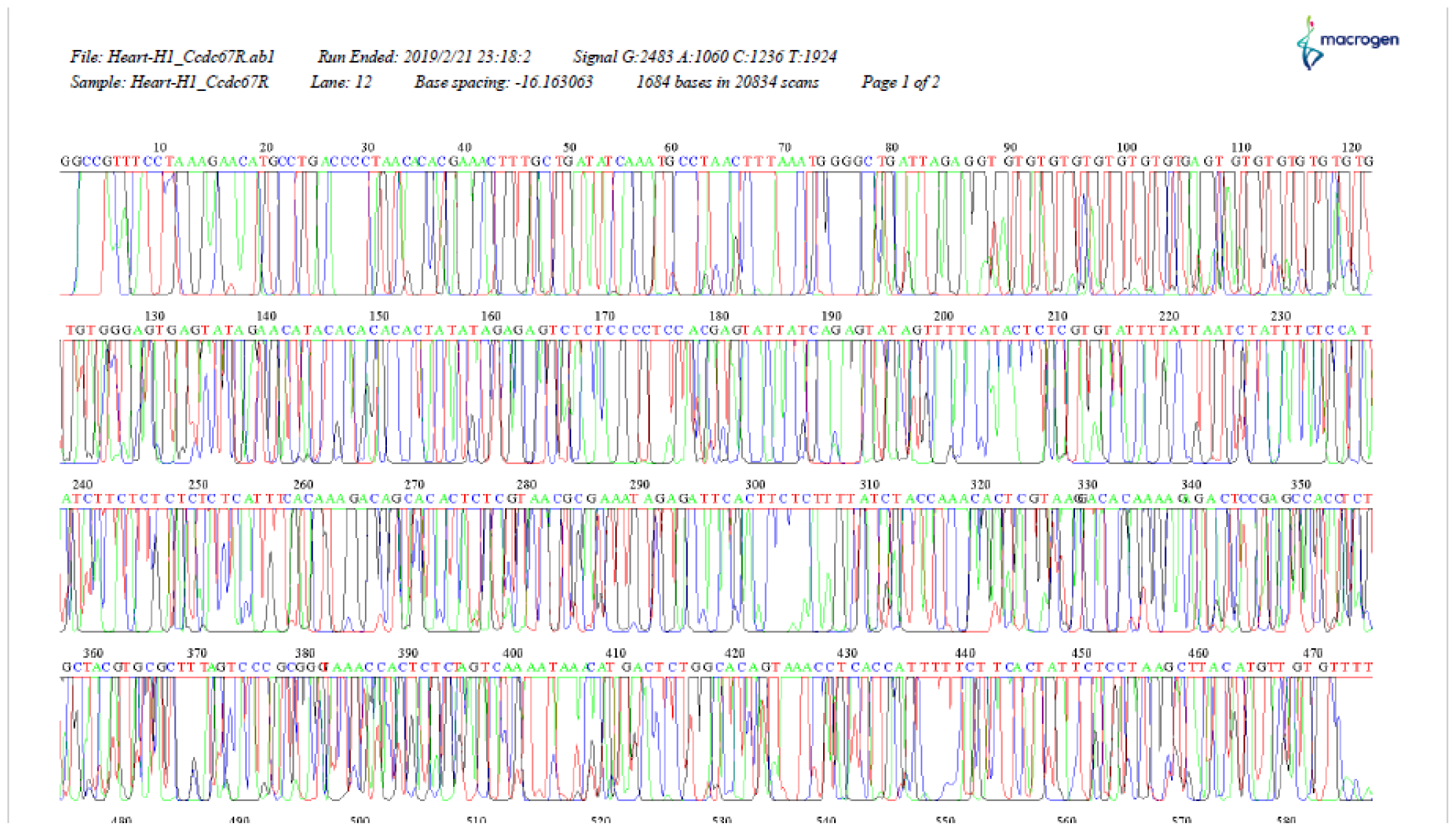

**>190221-R02_E03_Heart-H1_Ccdc67R.ab1 1684**

**GGCCGTTTCCTAAAGAACATGCCTGACCCCTAACACACGAAACTTTGCTGATATCAAATGCCTAACTTTAAATGGGGCTGATTAGAGGTGTGTGTGTGTGTGTGTGAGTGTGTGTGTGTGTGTGTGGGAGTGAGTATAGAACATACACACACACTATATAGAGAGTCTCTCCCCTCCACGAGTATTATCAGAGTATAGTTTTCATACTCTCGTGTATTTTATTAATCTATTTCTCCATATCTTCTCTCTCTCTCATTTCACAAAGACAGCACACTCTCGTAACGCGAAATAGAGATTCACTTCTCTTTTATCTACCAAACACTCGTAAGGACACAAAAGAGACTCCGAGCCACCTCTGCTACGTGCGCTTTAGTCCCGCGGGTAAACCACTCTCTAGTCAAAATAAACATGACTCTGGCACAGTAAACCTCACCATTTTTCTTCACTATTCTCCTAAGCTTACATGTTGTGTTTTTTACAGGCTGTTTCGGCGCGGTCTCTTCCCCTAATATACTTTAGCCTGTTCGATGCTTTATCTCCCTCTATCTGTGTCGTCTTTCTCTTCTCCCATTTATCGGGCGCTTATACTCGCTCTTTAACCCTCCTCTTCGCCTTTTTCCTTCCACTATTTCCACCTCCCATTCTTTTATCTACTACCCAAATTTACTTTCGCGTTTTAGATCGGGGTTAGATAAGGCTTTTTTTTGTTTTGTTTGTTCTCTTTTTGTCGGTTTTTCTCTTCTGCGGTTCTGTTCTTTGTTCCATCGTGTTCGTGTAGTCTTTCGTGTTTGGCTTTTTCCGTTACGGCCCTTGTTTTTTTTTTCTTTGTCTCTTTGACTCTCTTTTTTTTGATGTATTAGCTGTCTCGTGGTCCCTCCTCTTTTTCTCCGTCGTTTCTCTTTTTCCCCTCCCTCTTTCCTTTCTTCCCCCTTTTTTTTATCGGACATGGCCCGTTACCCTCCCGCCCTAACCTTTATTAATCTTACCATTCCCTCCCGATCATAAACACACCTAAATTATAGTCACGCCTATTTTATATTATCTCCCCCA**

**CCGTTCCCCCCAATTTACTTTTTCCCCACCTGCCAATACCCACCGAGGGGCACATTCGTTCCCTACTACATTACTACCTTTTTCCATACCCTAACCATACCCCCCGTTATTTATTAATACGTAGTAATTATTATTACTCTTAACATATCACCGTCACTCCTATACCAATTTCGCTCCACCATTTACGTAATACCAATACCTATACCCACACCCCCTACCCCCCGCGCGTATTACTAATTATTCTCGCTCCCTTCCACTTTTTTTATTTATACGCCGTTTTTTTTACCCCGCCCCCTCCCACCCCCCGTATAACACGACATATAGCCCCCCTTATATTTATATTATATTTTTATTTATTCCCTATTTTTTTTTTACTTCCACTGCAGCCCTTTTTCTTTTCTACTTTTTCCCCTACCCATCTTCCCATCCTTTTTTATTTTTTTTTAACTCACTCCATCCCCACACCCTTCTCCTCCTCTCTCCCTCATTTATTATATTCCCCCACTCATATTTTTAATTTTAATTATTCCTATTATATAACTCACAGCTTTCATTCCCCCCCAACTATTTAATTTAATTTTTTTTTTATCCCCTCATTCTCTTTATTTTCTTTTCTTATTATATTACGTTACAC**

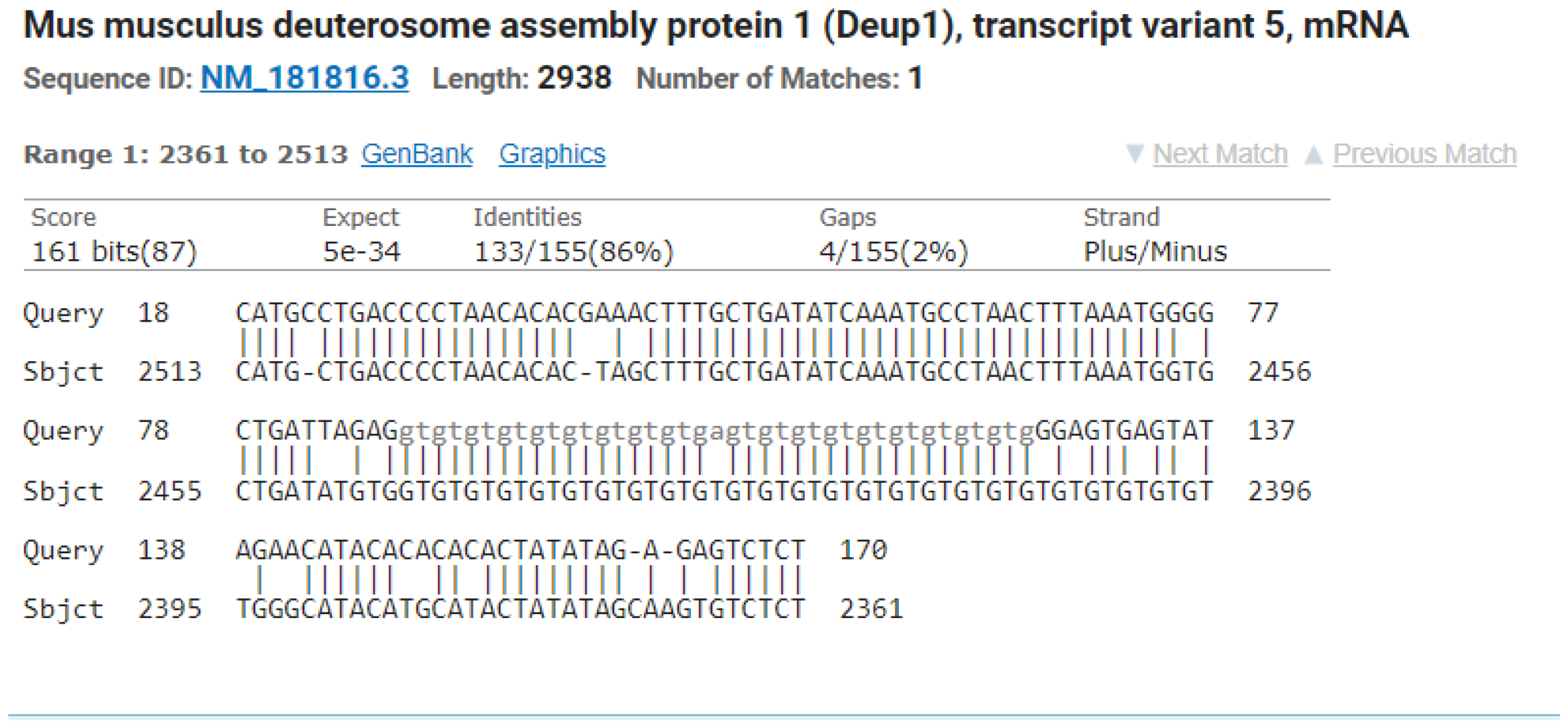

Both forward and reverse primers were sent with DNA for sequencing to macrogen, South Korea and for *CCDC67* it showed 93% identity to transcript variants 5,4,3,2 and 1. Only transcript variant 5 is shown for both F and R primers sequencing alignment.

## Notes

### Competing Interest Statement

The authors have declared no competing interest.

### Summary of Updates

The order of authors is revised.

